# Hydrogen sulfide improves the cold stress resistance through the CsARF5-CsDREB3 module in cucumber

**DOI:** 10.1101/2021.09.18.460893

**Authors:** Xiao-Wei Zhang, Xin Fu, Feng-Jiao Liu, Ya-Nan Wang, Huan-Gai Bi, Xi-Zhen Ai

## Abstract

Hydrogen sulfide (H_2_S) plays a crucial role in regulating cold tolerance. But the synergistic regulation of H_2_S and auxin in the plant response to cold stress has not been reported. In the study, we found that sodium hydrosulfide (NaHS, an H_2_S donor) treatment enhanced the cold tolerance of cucumber seedlings and increased the level of auxin. *CsARF5*, a cucumber auxin response factor (ARF) gene was isolated and its role in regulating H_2_S-mediated cold stress tolerance was described. Transgenic cucumber leaves overexpressing *CsARF5* were obtained. Physiological analysis indicated that overexpression of *CsARF5* enhanced the cold stress tolerance of cucumber and the regulation of the cold stress response by *CsARF5* depends on H_2_S. In addition, molecular assays showed that *CsARF5* modulated cold stress response by directly activating the expression of the dehydration-responsive element-binding (DREB)/C-repeat binding factor (CBF) gene *CsDREB3*, which was identified as a positive regulator of cold stress. Taken together, our results suggest that CsARF5 plays an important role in H_2_S-mediated cold stress in cucumber. These results shed light on the molecular mechanism by which H_2_S regulates cold stress response by mediating auxin signaling, and will provide insights for further studies on the molecular mechanism by which H_2_S regulates cold stress.

**Highlight:** Auxin signaling participates in H_2_S-mediated cold stress through the CsARF5-CsDREB3 module in cucumber.

## Introduction

Cucumber (*Cucumis sativus* L.) is one of the most important economic crops worldwide. The cultivation and yield of cucumbers in China have ranked among the top in the world for many years. In China, cucumber is generally grown in greenhouses. Because of the extreme low temperature conditions, the cucumber in greenhouse is prone to cold injury in winter and early spring. Cucumbers with cold injury showed inhibited growth, wilted and died in severe cases. Therefore, it is of great practical significance to study the effects of low temperature stress on cucumber growth and development and the response mechanism of cucumber to low temperature stress.

Numerous investigations have determined that hydrogen sulfide (H_2_S) is an important gas signaling molecule that plays a key role in the regulation of abiotic stress responses, including cold tolerance (Shi *et al*., 2015; Du *et al*., 2017; Hancock, 2019; Zhang *et al*., 2021). H_2_S is found to alleviate cold stress tolerance in many plant species, although the mechanisms remain elusive (Du *et al*., 2017; Fu *et al*., 2013; Shi *et al*., 2013; Nasibi *et al*., 2020). Several reports show that H_2_S modulates cold stress response possibly through mitogen activated protein kinase (MAPK) signaling (Du *et al*., 2017; Ba *et al*., 2021; Du *et al*., 2021). In addition to H_2_S, phytohormones, especially auxin, also play a vital positive regulatory role in cold stress response (Shibasaki *et al*., 2009; Rahman, 2013).

Auxin is involved in plant growth and development in various aspects, including cell division and elongation, tissue patterning, and the response to environmental stimuli (Zhao, 2010; Lv *et al*., 2019). Since auxin was identified for the first time in 1930s as indole-3-acetic acid, there has been a major breakthrough in the molecular mechanisms of auxin perception and signal transduction. Many genetic and biochemical approaches have elucidated that the TRANSPORT INHIBTOR RESPONSE1 (TIR1) protein functions as the receptor to perceive auxin signaling based on the reduced auxin response of *tir1* mutations (Dharmasiri *et al*., 2005; Kepinski and Leyser, 2005). Additionally, the SCF^TIR1^ ubiquitin-ligase complex is regarded as a central regulator of auxin signaling, and it-mediated proteolysis of auxin/indole acetic acid (Aux/IAA) proteins is responsible for auxin signaling transduction (Gray *et al*., 2001; Kepinski and Leyser, 2004). In this signaling pathway, Aux/IAA proteins are direct targets of TIR1. And Aux/IAA proteins directly interact with auxin response factors (ARFs) to repress their activities (Liscum and Reed, 2002). Upon exposure to auxin, the F-box protein TIR1 recruits the Aux/IAA proteins for degradation, which leads to the release of various auxin response factors, including *Small Auxin-up RNAs* (*SAURs*), *GH3s* and *Aux/IAAs*, and consequently regulates diverse auxin-mediated plant growth (Guilfoyle and Hagen, 2007; Weijers and Friml, 2009).

ARFs are vital transcription factors (TFs) that regulate the expression of auxin response genes (Guilfoyle and Hagen, 2007; Chandler *et al*., 2016; Li *et al*., 2016). To date, 23 and 25 *ARF* genes have been isolated in *Arabidopsis* and rice, respectively (Guilfoyle and Hagen, 2007; Wang *et al*., 2007; Shen *et al*., 2010). Most ARF members consist of a DNA-binding domain (DBD), a variable middle region and a carboxy-terminal dimerization domain (CTD) (Guilfoyle and Hagen, 2007; Tiwari *et al*., 2003). The DBD is classified as a plant-specific B3-type and functions to bind to TGTCTC/GAGACA sites (AuxREs) *in vitro* (Ulmasov *et al*., 1999; Tiwari *et al*., 2003). The middle region includes two types: activation domain-type (AD) and repression domain-type (RD), which are used as the basis of classification between transcription activators and transcription repressors (Tiwari *et al*., 2003; Li *et al*., 2016). Additionally, the CTD domain is responsible for protein-protein interactions by dimerizing with Aux/IAA proteins as well as other ARFs (Kim *et al*., 1997; Piya *et al*., 2014). Extensive studies have suggested that ARF proteins are involved in distinct developmental processes. In *Arabidopsis*, numerous ARF genes have been implicated in embryogenesis (ARF5 and ARF17) (Mallory *et al*., 2005), root growth (ARF7, ARF10, ARF16, and ARF19) (Okushima *et al*., 2005; Fukaki and Tasaka, 2009; Orosa-Puente *et al*., 2018; Lee *et al*., 2019), hypocotyl growth (ARF6, ARF7, ARF8, and ARF19) (Liu *et al*., 2018; Reed *et al*., 2018; Wang *et al*., 2020), shoot regeneration (ARF4 and ARF5) (Zhang *et al*., 2021), flower development (ARF2, ARF3, ARF6, and ARF8) (Nagpal *et al*., 2005; Finet *et al*., 2010), and senescence (ARF1 and ARF2) (Ellis *et al*., 2005). In the case of rice, genetic studies show that the functions of ARFs are different from the functions of *Arabidopsis*. OsARF1 is involved in root initiation and seed development (Inukai *et al*., 2005; Attia *et al*., 2009). OsARF12 regulates root elongation, iron accumulation, and phosphate homeostasis (Qi *et al*., 2012; Wang *et al*., 2014). OsARF16 regulates phosphate transport, phosphate starvation and iron deficiency responses (Shen *et al*., 2013; Shen *et al*., 2014; Shen *et al*., 2015). OsARF19 controls leaf angles (Zhang *et al*., 2015). OsARF11, OsARF12, OsARF16, and OsARF17 are involved in antiviral defence (Qin *et al*., 2020; Zhang *et al*., 2020). A recent study showed that OsARF6 regulates rice yield (Qiao et al., 2021). Though a number of ARF members have been functionally characterized in *Arabidopsis* and rice as mentioned earlier, little is known about the functions of *ARF* genes in cucumber.

In this study, we explored the molecular mechanisms by which H_2_S regulates cold stress response in cucumber. We found that H_2_S treatment could improve cold resistance and auxin content of cucumber. *CsARF5*, a transcriptional regulator in auxin signaling, was responsive to cold stress and H_2_S treatments, and overexpression of *CsARF5* improved the cold stress tolerance of cucumber. Further studies indicated that *CsARF5* modulated cold stress response by directly activating the expression of the dehydration-responsive element-binding (DREB)/C-repeat binding factor (CBF) gene *CsDREB3*.

## Materials and Methods

### Plant material and growth conditions

‘Jinyou 35’cucumber seedlings were used for cold stress treatment and genetic transformation. After soaking and germinating, the cucumber seeds were sown in nutrition bowls and transferred to a climate chamber with a PFD of 600 μmol m^−2^. s^−1^, a 25 °C/16 °C thermo-period, an 11-h photoperiod, and 80% relative humidity.

### Vector construction and transient transformation

To generate *CsARF5* and *CsDREB3* overexpression vectors, full-length *CsARF5* and *CsDREB3* were inserted into the pCAMBIA1300 plasmid.

For the transient transformation of detached cucumber leaves, cucumber leaves with the same growth conditions were taken and *Agrobacterium tumefaciens* LBA4404 (Weidi, Shanghai, China) with overexpression vector was injected into cucumber leaves through a medical syringe.

### Cold stress treatments

To evaluate the effect of H_2_S and IAA on the cold resistance of cucumber, cucumber seedlings with two-leaves were foliar sprayed with 1.0 mM Sodium hydrosulfide (NaHS, an H_2_S donor; Shanghai Macklin Biochemical Co., Ltd., Shanghai, China), 0.15 mM hypotaurine (HT, a specific scavenger of H_2_S; Sigma-Aldrich, Shanghai, China), and deionized water (H_2_O) respectively, or pretreated with 75 μM indole-3-acetic acid (IAA, Solarbio, Beijing, China), 50 μM 1-naphthylphthalamic acid (NPA, a polar transport inhibitor of IAA; Shanghai Aladdin Biochemical Technology Co., Ltd., Shanghai, China), or deionized water (H_2_O), respectively. 24 h later, half of the treatments were exposed to low temperatures (5 °C), and the other half of the seedlings were placed at normal temperatures as the control. The EL and accumulation of reactive oxygen species (ROS) were determined at 48 h after exposure of seedlings to 5 °C. Leaf samples of seedlings that pretreated with H_2_O and 1.0 mM NaHS were collected from 3 plants (n = 3) for transcriptome analysis after 6 h of cold stress.

To compare the difference in cold tolerance between WT and transgenic cucumber leaves, the pCAMBIA1300 empty vectors and overexpressed *CsARF5* and *CsDREB3* vectors were injected into the first leaf which was just flat of cucumber. 12 h later, WT leaves, *CsDREB3* overexpressing leaves, and some *CsARF5* overexpressing leaves were exposed to 5 °C. Other *CsARF5* overexpressing leaves were treated with HT and then exposed to 5 °C after the water droplets on the leaves were absorbed and dried. The gene expression of *CsDREB3*, *CsCBF1* and *CsCOR* in transgenic cucumber leaves was measured at 3 h after exposure to cold stress. NBT staining, DAB staining and ROS content were detected after cold treatment for 12 h.

### qRT-PCR analysis

The transcription levels of *CsARF5*, *CsDREB3*, *CsCBF1*, and *CsCOR* were examined using specific primers CsARF5 (qRT)-F/R, CsDREB3 (qRT)-F/R, CsCBF1 (qRT)-F/R, and CsCOR (qRT)-F/R, respectively. β-Actin was used as internal reference. All of the primers used are shown in Supplemental Table S1. qRT-PCR was carried out simultaneously with three biological replicates and three technical replicates.

### Detection of electrolyte leakage (EL)

EL was detected according to instructions described by Dong et al. (2013) (Dong *et al*., 2013). Leaf discs (0.2 g) were immersed in 20 ml deionized water and incubated at 25 °C for 3 h. The electrical conductivity (EC1) was estimated using a conductivity meter (DDB-303A, Shanghai, China). The leaf discs were boiled for 10 min, and then cooled to detect EC2. EL was calculated according to the following formula: EL= EC1/EC2 × 100.

### IAA content and flavin monooxygenase (FMO) activity assay

IAA content was measured using high-performance liquid chromatography-triple quadrupole mass spectrometry (HPLC–MS, Thermo Fisher Scientific, TSQ Quantum Access) according to the method of Li et al. (2014) with minor modifications by Zhang et al. (2020). In brief, leaf samples were extracted with 80% methanol (containing 30 μg.ml^−1^ sodium diethyldithiocarbamate) and the supernatant was retained rotary evaporation (Shanghai EYELA, N-1210B). Pigment and phenolic impurities of samples were removed using trichloromethane and polyvinylpolypyrrolidone (PVPP), respectively. Auxin was further extracted with ethyl acetate and then the ester phase was collected. Finally, rotation drying at 36 °C and drying were dissolved in 1.0 ml mobile phases (methanol: 0.04% acetic acid = 45:55, v/v). The filtrate can then be used directly for HPLC–MS analysis.

The activity of FMO was detected using an enzyme-linked immunosorbent assay (ELISA) kit (Jiangsu Meimian Industrial Co. Ltd, Jiangsu, China) as described by Zhang et al. (2020).

### Determination of H_2_O_2_ and superoxide anion (O_2_^.-^) contents

H_2_O_2_ content was estimated with the H_2_O_2_ kit (Nanjing Jiancheng Bioengineering Institute, Nanjing, China) according to the instructions. The O_2_^.-^ content was detected according to the method of Wang and Luo (1990). Cellular H_2_O_2_ and O_2_^.-^ were fluorescently stained with 2Ȳ, 7Ȳ-dichlorodihydrofluorescein diacetate (H_2_DCFDA, the fluorescent probe of H_2_O_2_) (MCE, Cat. No. HY-D0940, Shanghai, China) and dihydroethidium (DHE, O_2_^.-^ fluorescent probe) (Fluorescence Biotechnology Co. Ltd, Cat. No. 15200, Beijing, China), respectively as described by Galluzzi and Kroemer (2014) and modified by Zhang et al. (2020). In brief, the samples were infiltrated in 20 μM H_2_O_2_ fluorescent probe at 25 °C under dark conditions for 30 min. Then the tissues were rinsed with HEPES-NaOH buffer (pH 7.5). Under excitation at 488 nm and emission at 522 nm of inverted microscope (Leica DMi8), cellular H_2_O_2_ showed obvious green fluorescent coloration. For cellular O_2_^.-^ measurement, the samples were infiltrated in 10 μM DHE at 37 °C under dark conditions for 30 min. After fixation, the tissues were rinsed with Tris-HCl buffer (pH 7.5). O_2_^.-^ showed strong red fluorescence under excitation at 490 nm and emission at 520 nm under an inverted microscope (Leica DMi8).

### Nitroblue tetrazolium (NBT) and 3, 3-diaminobenzidine (DAB) staining

NBT staining of O_2_^.-^ was performed according to the method of Jabs et al. (1996) with minor modifications. The fresh leaves were washed with distilled water, immersed in 0.5 mM NBT in vacuum and stained at 28 °C for 1 h. Then the leaves were boiled in ethanol: lactic acid: glycerol (3:1:1) mixed solution to remove pigments, and O_2_^.-^ was visualized in blue-purple coloration. DAB staining of H_2_O_2_ was carried out as described by Thordal-Christensen et al. (1997). The cleaned fresh leaves were soaked in 1 mM DAB staining solution (pH 3.8) in vacuum and stained at 28 ◻ for 8 h. Then the leaves were boiled in ethanol: lactic acid: glycerol (3:1:1) mixed solution to remove pigments, and H_2_O_2_ was visualized in reddish-brown coloration.

### EMSAs

The CsARF5-HIS fusion protein and biotin labelled probes were prepared for EMSAs. The CsARF5-HIS fusion protein was obtained by inducing *Escherichia coli* BL21 (TransGen Biotech, Beijing, China) with isopropyl β-D-thiogalactoside (IPTG). The biotin-labelled probes were synthesized by Sangon Biotech (Shanghai) Co., Ltd. To perform the EMSAs, the fusion protein was mixed with the probe and incubated at 24 °C for 30 min. The protein-probe mixture was separated by nondenatured acrylamide gel electrophoresis.

### Dual luciferase assay

The promoter sequence of *CsDREB3* was amplified and cloned into pGreenII 0800-LUC to generate the reporter construct pCsDREB3-LUC. The effector plasmid was constructed by inserting full-length *CsARF5* into pGreenII 62-SK. Different plasmid combinations were injected into tobacco (*Nicotiana benthamiana*) leaves by *Agrobacterium tumefaciens* LBA4404. The leaves were sprayed with 100 mM luciferin and luminescence was detected after being placed in darkness for 3 min. Fluorescence images were obtained with a live imaging system (Xenogen, USA). The fluorescence activity was determined using a fluorescence activity detection kit (Promega, Madison, WI, USA).

### Statistical analysis

All experiments were performed at least in triplicate, and the results are expressed as the mean ± standard deviation (SD) of three replicates. The data were analyzed statistically with DPS software. Duncan’s multiple range test was used to compare differences among treatments, and *P* < 0.05 was considered statistically significant.

### Accession numbers

CsARF5 (CsaV3_3G045690), CsDREB3 (CsaV3_2G030880), CsCBF1 (XM_004140746), CsCOR (XM_011659051).

## Results

### NaHS improves cold tolerance in cucumber seedlings

In a previous study, we demonstrated that NaHS could improve the cold tolerance of cucumber seedlings in a concentration-dependent manner, and 1.0 mM NaHS treatment showed a very significant difference compared with the control (Zhang *et al*., 2020). Here, we found that 1.0 mM NaHS significantly reduced cold stress injury, accumulation of H_2_O_2_ and O_2_^.-^, as well as EL in cucumber seedlings after exposure to 5°C for 48 h. However, 0.15 mM H_2_S scavenger HT increased cold stress injury, H_2_O_2_, O_2_ ^.-^ and EL, compare with the deionized water (H_2_O, as a comparison)-treated seedlings (Fig. 1A–F). We also noticed that the mRNA abundance of *CsCBF1* and *CsCOR* in NaHS treated-seedlings increased by 1.38-fold and 3.14-fold, respectively under cold stress, but no obvious differences were observed between H_2_O and HT treatments (Fig. 1G, H). Therefore, we further confirmed that H_2_S improves cold tolerance in cucumber.

**Figure 1.**
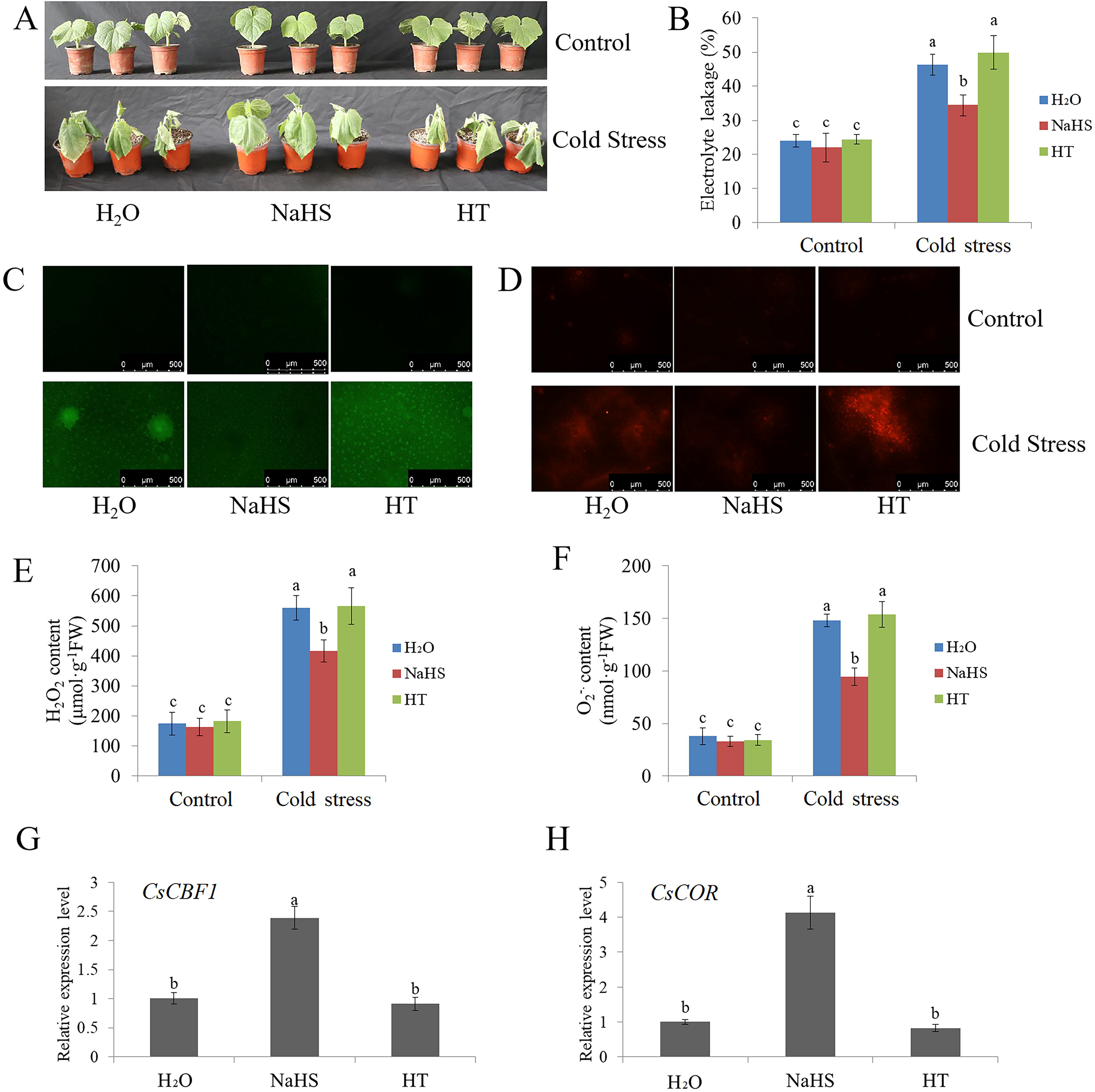
Effects of NaHS and HT on the cold tolerance of cucumber seedlings. **(A)** Phenotypic characterization of cucumber seedlings treated with NaHS or HT under cold stress (5 °C for 48 h). Water treatment was used as the control. H_2_O: water treatment; NaHS: NaHS (hydrogen sulfide provider) treatment; HT: HT (hydrogen sulfide scavenger) treatment. Control: cucumber seedlings before cold stress treatment; Cold stress: cucumber seedlings after cold stress treatment. Each treatment contained 5-10 cucumber seedlings. The experiments were repeated three times with similar results. A typical picture is shown here. **(B)** The EL results of cucumber seedlings before (control) and after cold stress for 48 h. **(C-D)** Inverted fluorescence microscopy imaging of H_2_O_2_ and O_2_^.-^ levels in cucumber seedling leaves before (control) and after cold stress treatment for 48 h. **(E-F)** Detection of H_2_O_2_ and O_2_^.-^ content of cucumber seedlings before (control) and after cold stress treatment for 48 h. **(G-H)** Expression of *CsCBF1* and *CsCOR* genes in cucumber seedlings treated with H_2_O, NaHS or HT under cold stress treatment for 6 h. qRT-PCR was performed simultaneously with three biological replicates and three technical replicates. The value of the water treatment was used as the reference and was set to 1. Error bars denote standard deviations. Different letters indicate significant differences (*P* < 0.05) based on Duncan’s multiple range tests.

### NaHS treatment affects auxin signaling

To explore the molecular mechanism by which H_2_S improves cold resistance in cucumber, we performed transcriptome analysis of cucumber seedlings treated with NaHS and H_2_O. A total of 1952 cucumber genes were analyzed from transcriptome data (Fig. 2A). Among these cucumber genes, 118 genes were downregulated and 54 genes were upregulated (Fig. 2A). We further analyzed the upregulated genes (Fig. 2B). One of the genes that caught our attention was an auxin response gene (accession number: CsaV3_3G045690, Fig. 2B). NCBI database comparison found that it was the *CsARF5* gene.

**Figure 2.**
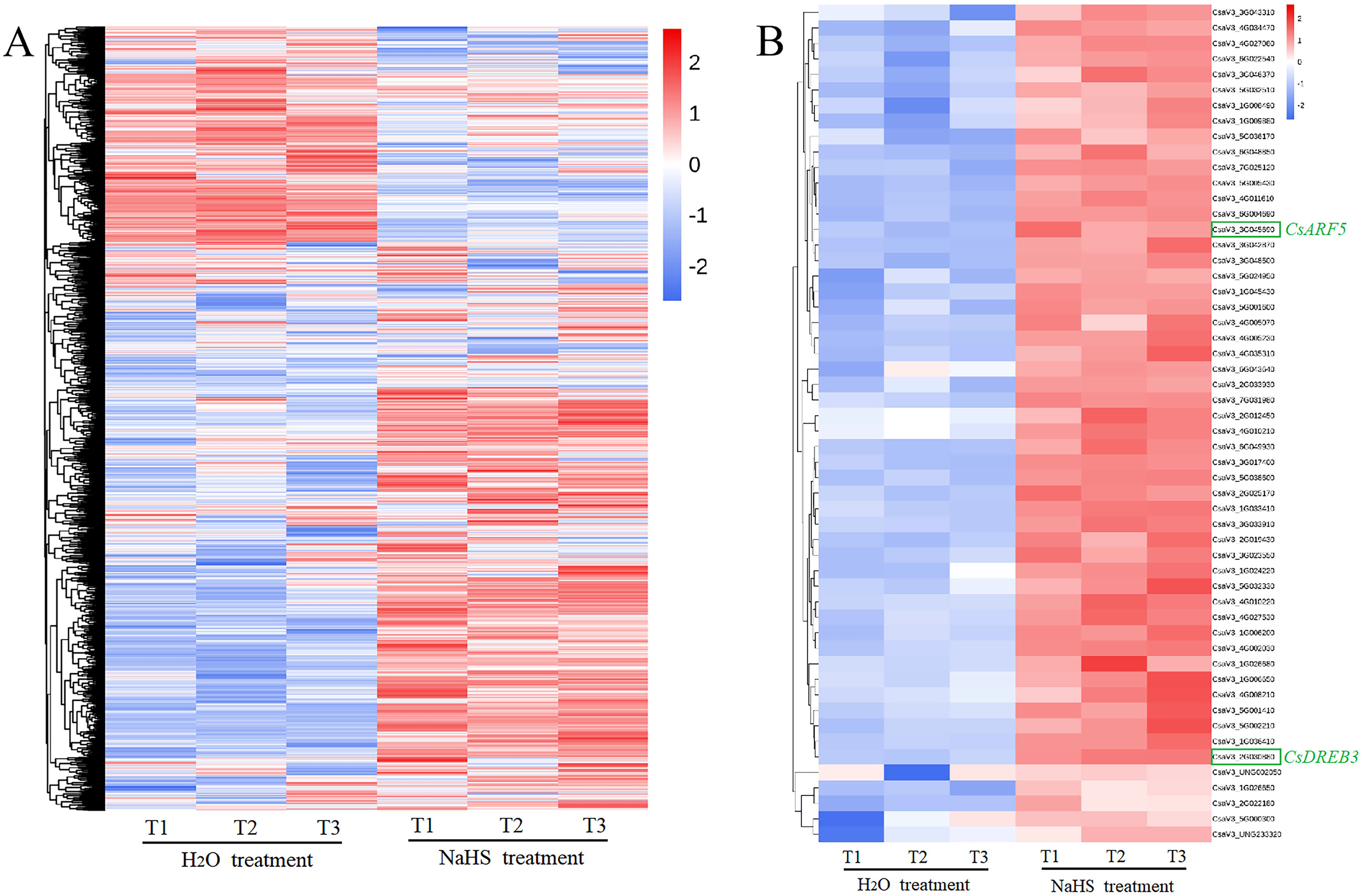
RNA- seq analysis of cucumber seedlings with or without NaHS treatment. **(A)** Normalized heat map showing the changes in gene expression after NaHS treatment at 5 °C for 6 h. All experiments were performed in triplicate. **(B)** Normalized heat map of upregulated gene expression after NaHS treatment at 5 °C for 6 h. All experiments were performed in triplicate.

To study the effect of NaHS on auxin signaling, we estimated the change in auxin content in cucumber seedlings treated with NaHS or HT. 1.0 mM NaHS markedly increased endogenous IAA accumulation and FMO (a key enzyme in auxin synthesis) activity. However, HT treatment revealed lower or similar IAA content and FMO activity compared with H_2_O treatment (Fig. 3A, B). In addition, NaHS treatment also significantly upregulated the mRNA levels of *CsARF5* and *CsDREB3*, while HT-treated seedlings showed no marked differences in the relative mRNA expression of *CsARF5* and *CsDREB3* relative to the H_2_O treatment (Fig. 3C, D). These data indicate that H_2_S affects auxin signaling in cucumber seedlings under cold-stress.

**Figure 3.**
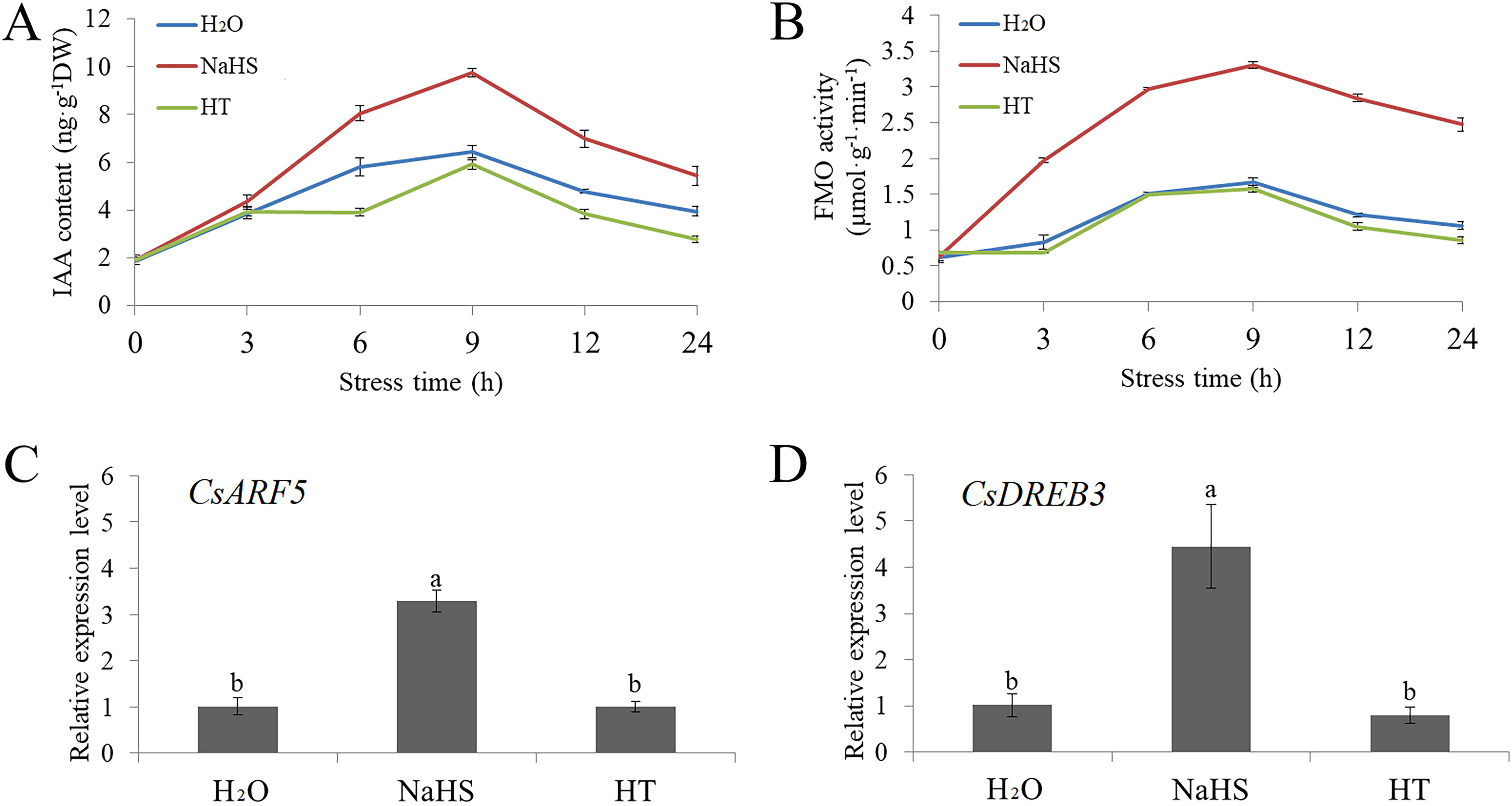
NaHS treatment affects IAA level in cucumber. **(A)** The IAA content of cucumber seedlings treated with NaHS or HT under cold stress treatment for 24 h. **(B)** The FMO activity of cucumber seedlings treated with NaHS or HT under cold stress treatment for 24 h. **(C)** Expression of the *CsARF5* gene in cucumber seedlings treated with NaHS or HT under cold stress treatment for 6 h. **(D)** Expression of the *CsDREB3* gene in cucumber seedlings treated with NaHS or HT under cold stress treatment for 6 h. qRT-PCR was performed simultaneously with three biological replicates and three technical replicates. The value of 0 h was used as the reference and was set to 1. Error bars denote standard deviations. Different letters indicate significant differences (*P* < 0.05) based on Duncan’s multiple range tests.

### IAA treatment improves cold resistance of cucumber seedlings

Our previous study proved that 75 μM IAA enhances the cold tolerance of cucumber seedlings (Zhang *et al*., 2020). Here, we found that 75 μM IAA significantly reduced the EL, and accumulation of H_2_O_2_ and O_2_^.-^ caused by cold stress, while 50 μM 1-naphthylphthalamic acid (NPA, a polar transport inhibitor) treatment showed no remarkable difference relative to H_2_O treatment under cold stress (Fig. 4A–F). As an auxin response factor, the relative mRNA expression of *CsARF5* was up-regulated in IAA-treated seedlings under cold stress. However, no remarkable difference was found in the mRNA expression of *CsARF5* between the NPA and H_2_O treatments (Fig. 4G). The mRNA expression levels of *CsCBF1* and *CsCOR* were significantly upregulated in IAA-treated seedlings, but downregulated or not influenced in HT-treated seedlings compared with H_2_O-treated seedlings when exposed to cold stress (Fig. 4H, I). The latest results are in keeping with our earlier findings, so we further confirm that IAA enhances cold tolerance in cucumber.

**Figure 4.**
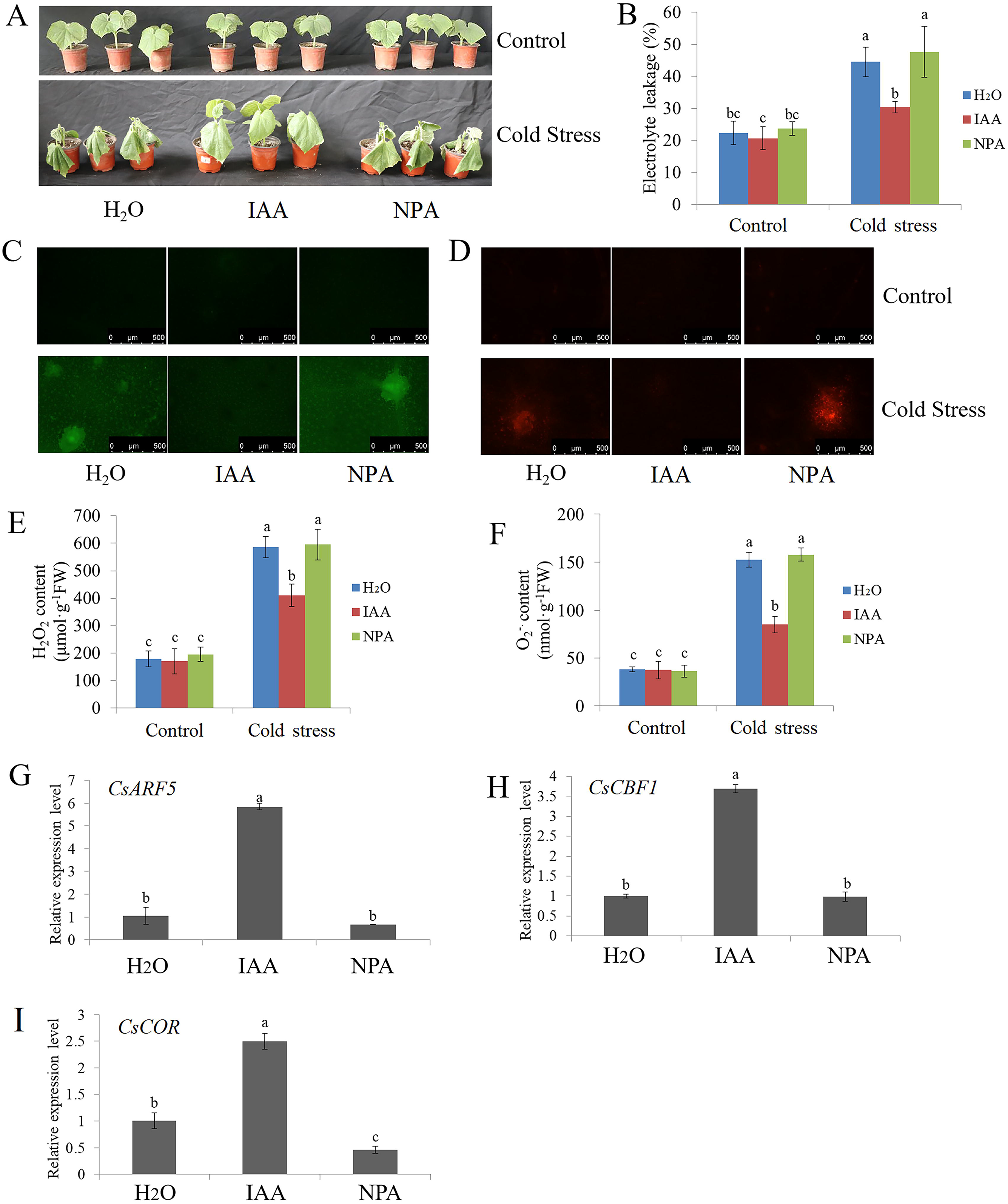
Effects of IAA and NPA on the cold resistance of cucumber seedlings. **(A)** Phenotypic characterization of cucumber seedlings treated with IAA or NPA under cold stress (5 °C for 48 h). Water treatment was used as the control. H_2_O: water treatment; IAA: indole-3-acetic acid treatment; NPA: NPA (a polar transport inhibitor) treatment. Control: cucumber seedlings before cold stress treatment; Cold stress: cucumber seedlings after cold stress treatment. Each treatment contained 5-10 cucumber seedlings. The experiments were repeated three times with similar results. A typical picture is shown here. **(B)** The EL results of cucumber seedlings before (control) and after cold stress treatment for 48 h. **(C-D)** Inverted fluorescence microscope imaging of H_2_O_2_ and O_2_^.-^ levels in cucumber seedling leaves before (control) and after cold stress treatment for 48 h. **(E-F)** Detection of H_2_O_2_ and O_2_^.-^ accumulation of cucumber seedlings before (control) and after cold stress treatment for 48 h. **(G-I)** Expression of *CsARF5*, *CsCBF1* and *CsCOR* genes in cucumber seedlings treated with H_2_O, IAA or NPA under cold stress for 6 h. qRT-PCR was performed simultaneously with three biological replicates and three technical replicates. The value of the water treatment was used as the reference and was set to 1. Error bars denote standard deviations. Different letters indicate significant differences (*P* < 0.05) based on Duncan’s multiple range tests.

### CsARF5 positively regulates cold stress tolerance of cucumber

We wondered whether the auxin response gene *CsARF5* is involved in the H_2_S-mediated cold stress response. qRT-PCR results showed that cold stress increased the mRNA abundance of the *CsARF5* gene, and the expression reached a peak after seedlings were exposed to cold for 3 h, and then decreased gradually (Fig. 5A). Compared with the control and HT treatments, NaHS treatment further increased the expression of *CsARF5* (Fig. 5B). These results demonstrate that *CsARF5* is responsive to cold stress and H_2_S treatment.

**Figure 5.**
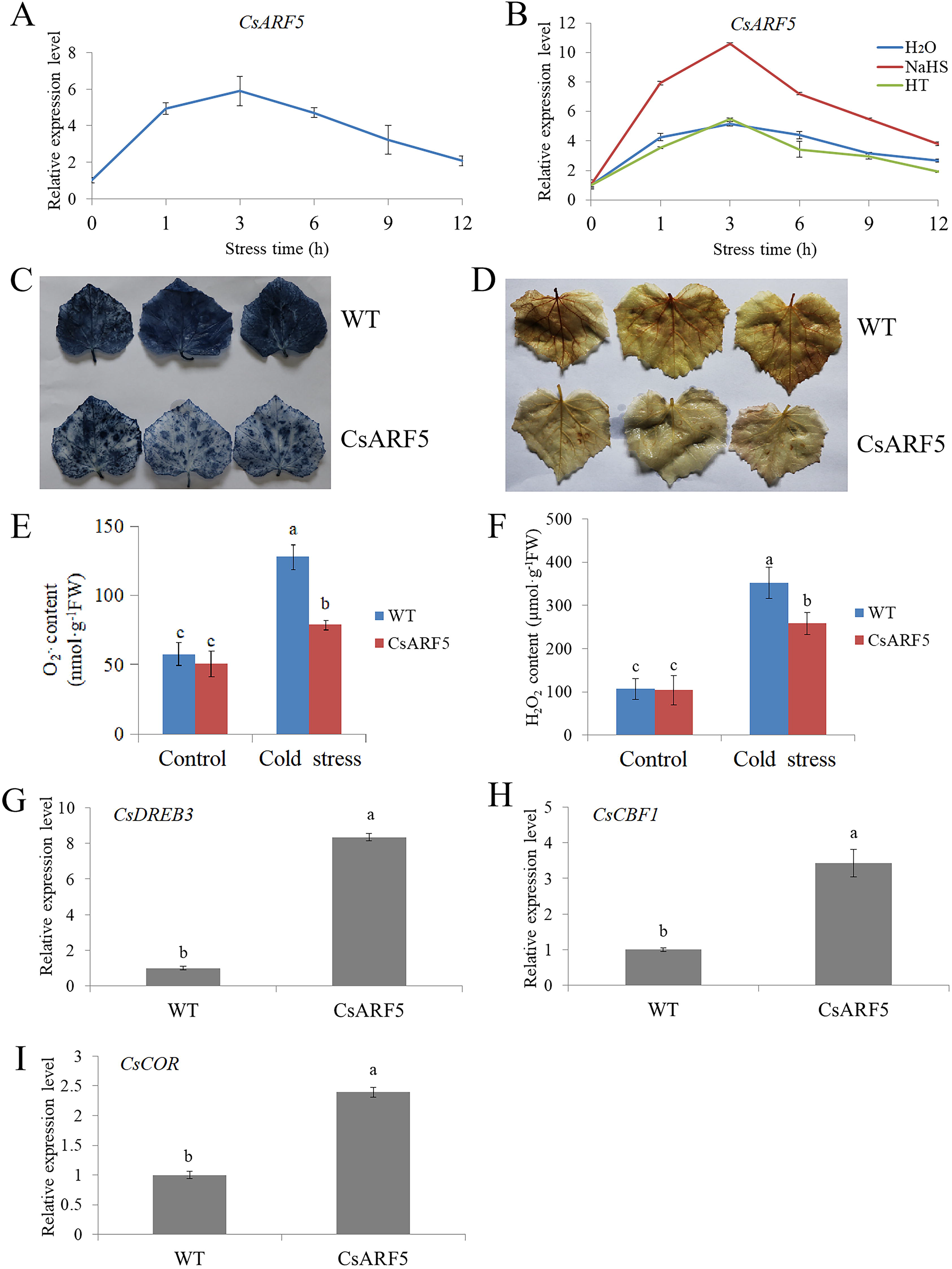
Overexpression of *CsARF5* improves cold tolerance of cucumber. **(A)** Expression of the *CsARF5* gene in cucumber seedlings under cold stress treatment for 12 h. **(B)** Expression of the *CsARF5* gene in cucumber seedlings treated with NaHS or HT under cold stress treatment for 12 h. **(C-D)** NBT and DAB staining of empty vector control (WT) and *CsARF5* transient transgenic cucumber leaves treated with cold stress for 12 h. Each genotype contained 5-10 cucumber leaves. The experiments were repeated three times with similar results. A typical picture is shown here. **(E-F)** Detection of O_2_^.-^ and H_2_O_2_ contents of transgenic cucumber leaves before (control) and after cold stress treatment for 12 h. **(G-I)** Expression of *CsDREB3*, *CsCBF1* and *CsCOR* genes in transgenic cucumber leaves under cold stress for 3 h. qRT-PCR was performed simultaneously with three biological replicates and three technical replicates. The value of WT was used as the reference and was set to 1. Error bars denote standard deviations. Different letters indicate significant differences (*P* < 0.05) based on Duncan’s multiple range tests.

To further explore the role of *CsARF5* in the response to cold stress in cucumber, we obtained *CsARF5* transgenic cucumber leaves through *Agrobacterium*-mediated transient genetic transformation (Supplemental Fig. 1A). Then, we observed the accumulation of ROS in transgenic leaves after exposure to cold stress for 12 h, using NBT and DAB staining. The results showed that the accumulation of ROS in cucumber leaves overexpressing with *CsARF5* (CsARF5) was less than that of the empty vector control (WT) under cold stress (Fig. 5C–D). In addition, the contents of O_2_ ^.-^ and H_2_O_2_ were measured with biochemical analysis, and the results were in agreement with the NBT and DAB staining images (Fig. 5E–F). qRT-PCR results revealed that *CsARF5* overexpression upregulated the mRNA level of cold stress-responsive genes *CsDREB3*, *CsCBF1*, and *CsCOR* (Fig. 5G–I). These data suggest that *CsARF5* is a positive regulator of cold stress response.

### HT treatment affects CsARF5-mediated cold stress tolerance

Considering that H_2_S induces the expression of *CsARF5*, and that *CsARF5* positively regulates cold stress resistance, we wondered about the role of *CsARF5* in H_2_S-mediated cold stress. The H_2_S scavenger HT was applied to *CsARF5*-overexpressing cucumber leaves to observe ROS accumulation. The NBT and DAB staining results showed that the application of HT alleviated the CsARF5-decreased ROS accumulation in detached cucumber leaves (Fig. 6A–B). The biochemical analysis for O_2_^.-^ and H_2_O_2_ was consistent with the NBT and DAB staining results (Fig. 6C–D). qRT-PCR results showed that the application of HT inhibited the promotion of *CsDREB3*, *CsCBF1*, and *CsCOR* mRNA expression levels caused by CsARF5 (Fig. 6E–G). These results suggest that the regulation of the cold stress response by *CsARF5* depends on H_2_S.

**Figure 6.**
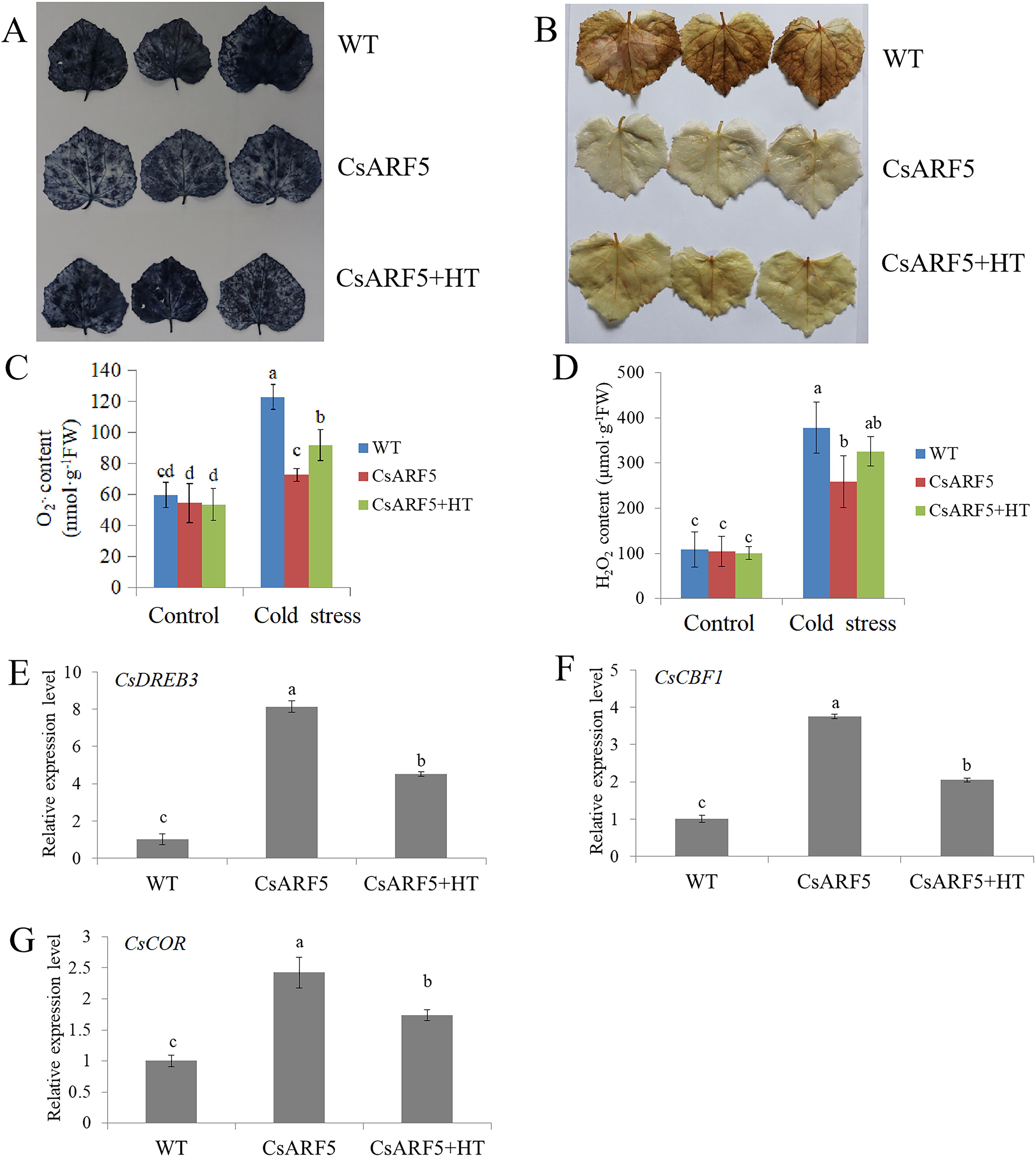
Effect of HT on the cold resistance of *CsARF5* transgenic cucumber. **(A-B)** NBT and DAB staining of empty vector control (WT), *CsARF5* transient transgenic cucumber leaves (CsARF5) and CsARF5 sprayed with HT (CsARF5+HT) treated with cold stress for 12 h. Each genotype contained 5-10 cucumber leaves. The experiments were repeated three times with similar results. A typical picture is shown here. **(C-D)** Detection of O ^.-^ and H O contents of transgenic cucumber leaves before (control) and after cold stress or HT treatment for 12 h. **(E-G)** Expression of *CsDREB3*, *CsCBF1* and *CsCOR* genes in transgenic cucumber leaves under cold stress for 3 h. qRT-PCR was performed simultaneously with three biological replicates and three technical replicates. The value of WT was used as the reference and was set to 1. Error bars denote standard deviations. Different letters indicate significant differences (*P* < 0.05) based on Duncan’s multiple range tests.

### Overexpression of CsDREB3 enhances cold stress tolerance of cucumber

DREB/CBF TFs play essential roles in the regulation of the plant cold stress response (Zhou *et al*., 2010; Lata and Prasad, 2011). *CsDREB3* was induced by NaHS (Fig. 2B), which prompted us to explore whether CsDREB3 was involved in the cold stress response in cucumber. As shown in Fig. 7A, cold stress induced the expression of the *CsDREB3* gene, and the expression reached a peak at 3 h, and then decreased gradually. Compared with the control and HT treatments, NaHS treatment further increased the expression of *CsDREB3* (Fig. 7B).

**Figure 7.**
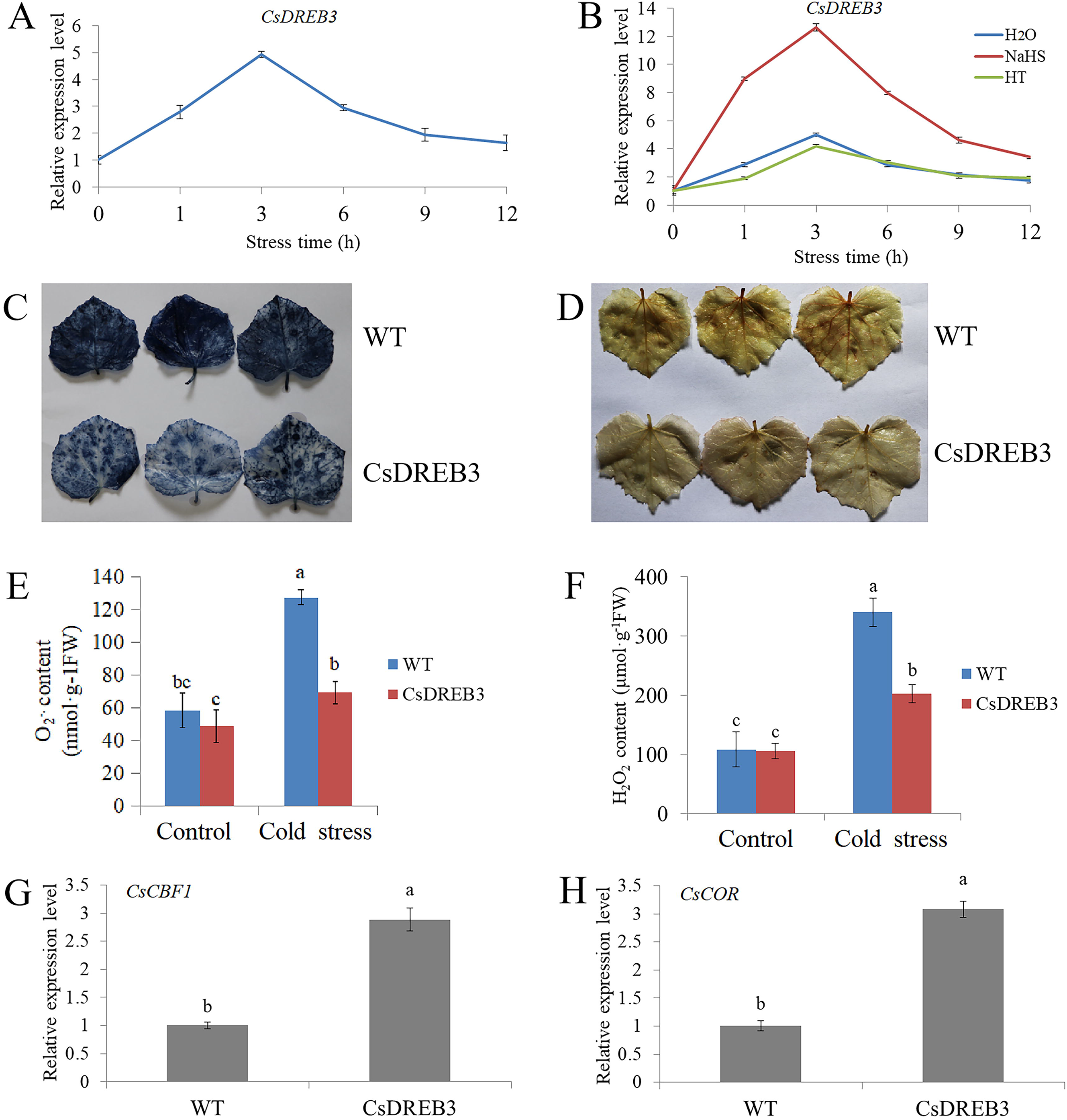
Overexpression of *CsDREB3* improves the cold tolerance of cucumber. (A) Expression of the *CsDREB3* gene in cucumber seedlings under cold stress treatment for 12 h. Expression of the *CsDREB3* gene in cucumber seedlings treated with NaHS or HT under cold stress treatment for 12 h. **(C-D)** NBT and DAB staining of empty vector control (WT) and *CsDREB3* transient transgenic cucumber leaves treated with cold stress for 12 h. Each genotype contained 5-10 cucumber leaves. The experiments were repeated three times with similar results. A typical picture is shown here. **(E-F)** Detection of O_2_^.-^ and H_2_O_2_ contents of transgenic cucumber leaves before (control) and after cold stress treatment for 12 h. **(G-H)** Expression of *CsCBF1* and *CsCOR* genes in transgenic cucumber leaves under cold stress for 3 h. qRT-PCR was performed simultaneously with three biological replicates and three technical replicates. The value of WT was used as the reference and was set to 1. Error bars denote standard deviations. Different letters indicate significant differences (*P* < 0.05) based on Duncan’s multiple range tests.

To investigate the role of *CsDREB3* in the response to cold stress in cucumber, *CsDREB3* transient transgenic cucumber leaves were obtained (Supplemental Fig. 1B). NBT and DAB staining results displayed that overexpression of *CsDREB3* decreased ROS accumulation after cold stress treatment (Fig. 7C–D). In addition, O_2_^.-^ and H_2_O_2_ detection results also revealed that the accumulation of O_2_^.-^ and H_2_O_2_ was significantly lower in leaves of overexpressioning *CsDREB3* than in WT leaves (Fig. 7E–F). qRT-PCR results suggested that overexpression of *CsDREB3* increased the expression of *CsCBF1* and *CsCOR* (Fig. 7G–H). These data demonstrate that *CsDREB3* positively regulates the cold tolerance of cucumber.

### CsARF5 directly activates the expression of CsDREB3

CsARF5 acts as an auxin response factor and can bind to the AuxRE motif in the promoters of target genes (Guilfoyle and Hagen, 2007). Considering the similar expression patterns of CsARF5 and CsDREB3 in cold stress and H_2_S treatment, as well as the key role of DREB/CBF TFs in the regulation of the cold stress response, we hypothesized that CsARF5 might be involved in the response to cold stress by mediating the expression of CsDREB3. Then, we analyzed the sequence of the *CsDREB3* gene promoter region and found a putative AuxRE motif (Fig. 8A). Fortunately, the directly binding between CsARF5 protein and the promoter of *CsDREB3* was detected by electromobility shift assay (EMSA) (Fig. 8B). To test how CsARF5 regulated the expression of *CsDREB3*, dual luciferase assays in tobacco leaves were performed. The CsARF5 effector construct was expressed under the 35S promoter and the promoter of *CsDREB3* was fused to the Luc gene as a reporter (Fig. 8C). The results showed that co-expression of 35Spro:CsARF5 with CsDREB3pro: Luc led to an obvious increase in luminescence intensity (Fig. 8D–E), while the binding site was mutated, and the activation was abolished (Fig. 8D–E). These results suggest that CsARF5 trans-activates the expression of *CsDREB3* in cucumber.

**Figure 8.**
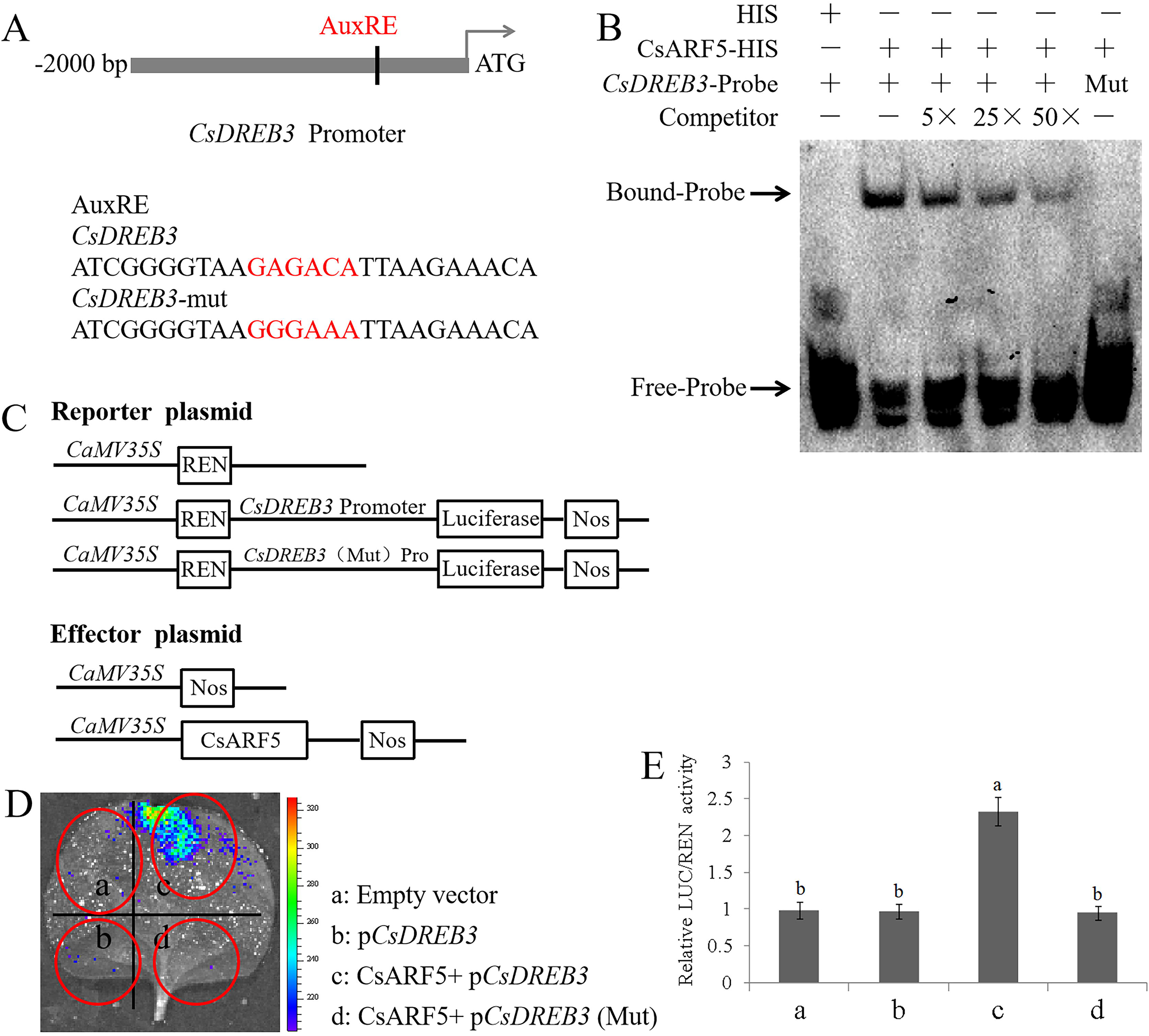
CsARF5 directly modulates the expression of *CsDREB3*. **(A)** Schematic diagram showing the *CsDREB3* promoter probe used for EMSAs. AuxRE indicates the CsARF5 binding site. Mutated probe (*CsDREB3-*mut) in which the 5’-GAGACA-3’ motif was replaced by 5’-GGGAAA-3’. **(B)** EMSAs showing that CsARF5 binds to the *MdDREB3* promoter. “–” indicates the absence of corresponding proteins or probes. “+” indicates the presence of corresponding proteins or probes. “5×”, “25×” and “50×” indicate increased probe concentrations. The experiments were repeated three times with similar results. A typical picture is shown here. **(C)** Schematic representation of the LUC reporter vector containing the *CsDREB3* promoter and the effector vectors expressing *CsARF5* under the control of the 35S promoter. **(D)** Dual luciferase tests in tobacco leaves showing that CsARF5 activates *CsDREB3* transcription. Empty vector: pGreenII 62-SK + pGreenII 0800-LUC; pCsDREB3: pGreenII 62-SK + pCsDREB3-pGreenII 0800-LUC; CsARF5 + pCsDREB3: CsARF5-pGreenII 62-SK + pCsDREB3-pGreenII 0800-LUC; CsARF5 + pCsDREB3(Mut): CsARF5-pGreenII 62-SK + pCsDREB3(Mut)-pGreenII 0800-LUC. Mutated promoter sequence (pCsDREB3*-*Mut) in which the 5’-GAGACA-3’ motif was replaced by 5’-GGGAAA-3’. The experiments were repeated three times with similar results. A typical picture is shown here. **(E)** LUC/REN activity detection to verify that CsARF5 activates the transcription of *CsDREB3*. Empty vector was used as the reference and set to 1. Error bars denote standard deviations. Different letters indicate significant differences (*P* < 0.05) based on Duncan’s multiple range tests. All experiments were performed three times with similar results, and representative data from one repetition are shown.

## Discussion

H_2_S, as a gaseous signaling molecule, plays a crucial role in plant relevance to various stress conditions, such as low temperature, salt, drought, and heavy metals (Huang et al., 2021; Zhang et al., 2021). Exogenous application of the H_2_S donor NaHS can effectively improve plant growth and stress response (Guo et al., 2016). In this study, we found that NaHS treatment improved cold-tolerance of cucumber seedlings (Fig. 1), suggesting that the response of H_2_S to cold stress is consistent in different species. H_2_S and phytohormones have synergistic effects on the H_2_S-mediated plant stress response. For example, H_2_S interacts with ABA and ethylene and is involved in the plant response to drought stress and in the regulation of stomatal closure (Jin *et al*., 2013; Hou *et al*., 2013). SA may play a key role in H_2_S-alleviated heavy metal stress and low temperature stress (Li *et al*., 2015; Qiao *et al*., 2015). Here, we found that NaHS treatment promoted IAA synthesis and upregulated the expression of auxin-responsive genes (Fig. 2 and Fig. 3), indicating that H_2_S may crosstalk with auxin to regulate the cold stress response.

Several plant hormones, such as jasmonic acid, salicylic acid, and ethylene, have been shown to play key roles in the response to cold stress (Miura and Tada, 2014; Kazan, 2015; Hu *et al*., 2017). However, the role of auxin under cold stress is limited. Auxin is a crucial phytohormone that is involved in a variety of plant physiological and developmental processes including the regulation of the cold stress response (Zhao, 2018; Blakeslee *et al*., 2019). Previous investigations have determined that cold stress promotes auxin biosynthesis or changes auxin gradient distribution, thus affecting the root gravity response in *Arabidopsis*, rice, and poplar (Shibasaki *et al*., 2009; Popko *et al*., 2010; Du *et al*., 2013; Rahman, 2013). However, there are few studies on the effects of auxin to the cold stress response of plants. In this study, we found that the application of auxin improved cold resistance, whereas the application of the polar transport inhibitor NPA slightly reduced the cold resistance of cucumber seedlings (Fig. 4), demonstrating that auxin is a positive regulator of the cold stress response.

As a master regulator of auxin signaling, ARF TFs have been functionally characterized in *Arabidopsis* and rice (Guilfoyle and Hagen, 2007; Wang *et al*., 2007; Shen *et al*., 2010). An increasing number of studies have indicated that ARFs regulate multiple plant developmental processes such as root growth (Orosa-Puente *et al*., 2018; Lee *et al*., 2019), flower development (Nagpal *et al*., 2005; Finet *et al*., 2010), senescence (Ellis *et al*., 2005), and stress response (He *et al*., 2005; Rahman, 2013). Among these processes, the auxin-regulated cold stress response has become a major focus of biotechnology in cucumber with the task of identifying the cold resistance genes and improving the yield of cucumber. In this study, an ARF TF, CsARF5, was isolated from cucumber and its expression was induced by NaHS and cold stress treatments, suggesting that CsARF5 may be involved in H_2_S-mediated cold tolerance (Fig. 5A–B).

To investigate the functions of CsARF5, transgenic cucumber leaves overexpressing *CsARF5* were generated. As hypothesized, the physiology and genetic analysis indicated that the overexpression of *CsARF5* decreased the accumulation of ROS and increased the expression level of cold stress-responsive genes, thus improving the cold stress resistance of cucumber (Fig. 5). These results suggest that CsARF5 positively regulates the cold stress tolerance of cucumber. Meanwhile, the H_2_S scavenger HT was applied to *CsARF5*-overexpressing cucumber leaves to study the role of *CsARF5* in H_2_S-mediated cold stress. The results showed that HT inhibited the cold stress resistance increased by CsARF5 (Fig. 6), indicating that the regulation of the cold stress response by CsARF5 depends on H_2_S.

DREB/CBF proteins play important roles in the regulation of the plant cold stress response (Zhou *et al*., 2010; Lata and Prasad, 2011). In *Arabidopsis*, DREB1A, DREB2C, CBF1, CBF2, and CBF3 are involved in the cold stress response (Liu *et al*., 1998; Gilmour *et al*., 1998; Lee *et al*., 2010; Lata and Prasad, 2011). In rice, OsDREB1A, OsDREB1B, OsDREB1C, OsDREB1F, OsDREB2B, and OsDREBL are responsive to cold stress treatment (Chen *et al*., 2003; Dubouzet *et al*., 2003; Wang *et al*., 2008; Matsukura *et al*., 2010). Here, a cucumber *DREB* genes, namely *CsDREB3*, were induced by NaHS and cold stress treatments, and overexpression of *CsDREB3* significantly enhanced the cold stress tolerance of cucumber (Fig. 7), revealing that CsDREB3 is a positive regulator of cold stress.

ARF5 is known to function as a transcriptional activator (Guilfoyle and Hagen, 2007). The remarkable expression of *CsDREB3* in CsARF5 transgenic leaves prompted us to consider whether CsARF5 can directly regulate CsDREB3 expression (Fig. 5G). The putative AuxRE elements that were recognized by CsARF5 were searched in the promoter region of CsDREB3. Fortunately, one AuxRE site was found, and the results of EMSAs and dual luciferase assays provided evidence to show that CsARF5 could bind to the promoter of *CsDREB3* and activate its expression (Fig. 8).

Based on previous studies in model plant species and our results in this study, a working model of CsARF5 regulating the H_2_S-mediated cold stress response was proposed (Fig. 9). CsIAA proteins interact with CsARF5 and interfere with the transcriptional regulation of *CsDREB3* by CsARF5. When H_2_S signaling was detected, auxin levels in cucumber were increased. CsARF5 is freed from the CsIAA protein complex. Then, CsARF5 specifically activated the expression of *CsDREB3* to improve the cold tolerance of cucumber. Our study elucidates the molecular mechanism by which H_2_S regulates the cold stress response in cucumber by mediating auxin signaling, which will provide insights for further studies on the molecular mechanisms by which H_2_S regulates cold stress. A better understanding of the function and signal transduction of CsARF5 in cucumber is helpful to regulate the resistance of cucumber to low temperature stress, to obtain high-quality fruit.

**Figure 9.**
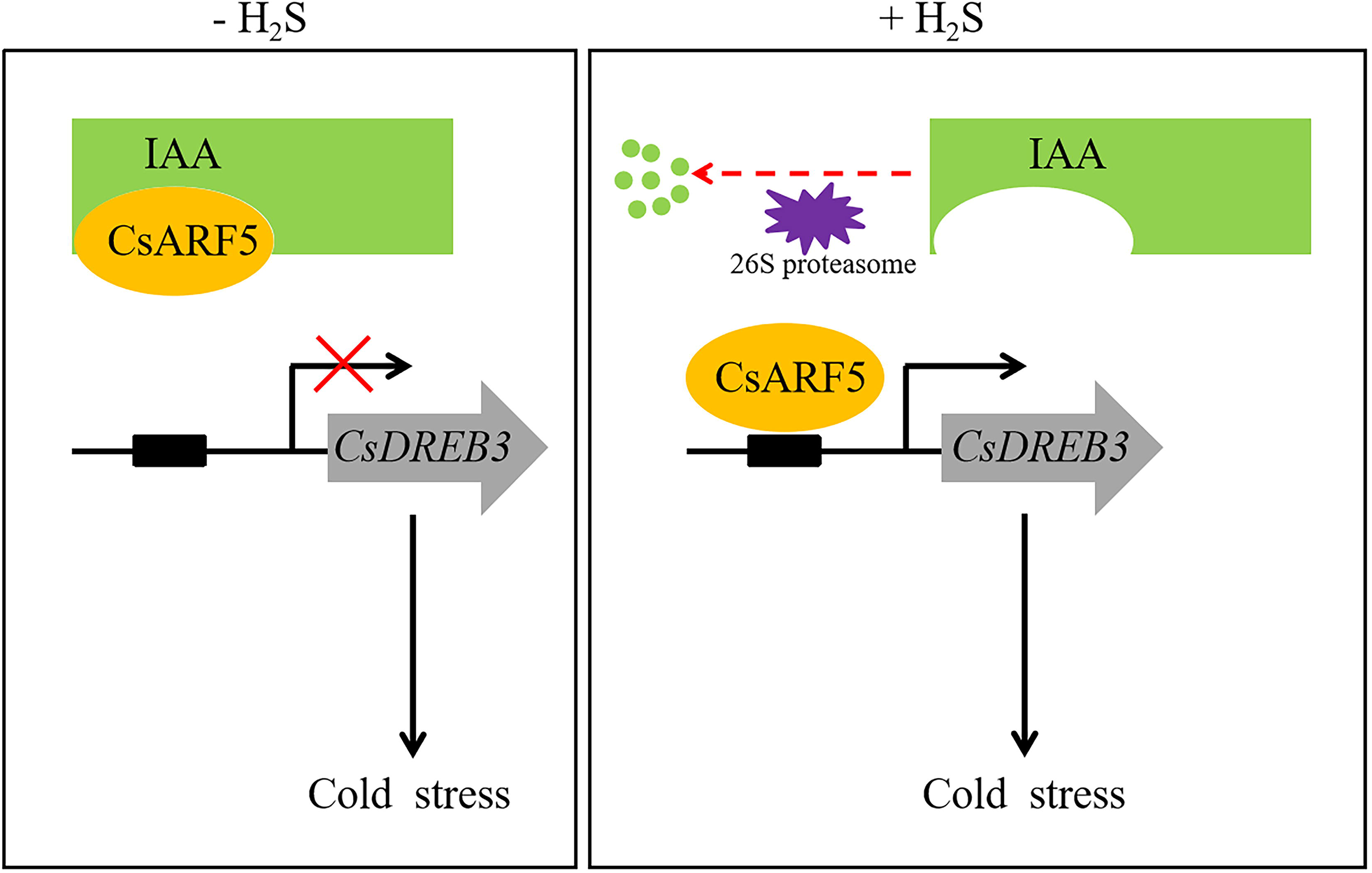
A regulatory model elucidates that H_2_S mediates the cold stress response in cucumber through auxin signaling. In the absence of H_2_S, IAA repressor proteins inhibit the expression of CsARF5, which in turn inhibits the cold stress response mediated by the CsARF5-*CsDREB3* module. In the presence of H_2_S, IAA repressor proteins release CsARF5, which promotes the cold stress response by activating *CsDREB3* expression.

## Data Availability

All data generated or analyzed during this study are included in this published article.

## Acknowledgements

This work was supported by grants from The National Science Foundation of China (31572170), The National Key Research and Development Program of China (2018YFD1000800), The Major Science and Technology Innovation of Shandong Province in China (2019JZZY010715), and The Special Fund of Vegetable Industrial Technology System of Shandong Province (SDAIT-05–10).

## Author contributions

Xi-Zhen Ai and Xiao-Wei Zhang designed the experiment. Xiao-Wei Zhang performed the research and analyzed the data. Xin Fu, Feng-Jiao Liu, Ya-Nan Wang and Huan-Gai Bi worked together with Xiao-Wei Zhang to accomplish the experiment. Xi-Zhen Ai and Xiao-Wei Zhang wrote the paper.

## Supplemental data

**Supplemental Tables**

**Supplemental Table 1.** Primers used for gene expression analysis and vector construction.

**Supplemental Table 2.** The promoter sequence of CsDREB3.

**Supplemental Figures**

**Supplemental Figure 1.** Identification of transient transgenic cucumber leaves.

